# The antibiotic bedaquiline activates host macrophage innate immune resistance to bacterial infection

**DOI:** 10.1101/758318

**Authors:** Alexandre Giraud-Gatineau, Juan Manuel Coya, Alexandra Maure, Anne Biton, Michael Thomson, Elliott M. Bernard, Maximiliano G. Gutierrez, Gérald Larrouy-Maumus, Roland Brosch, Brigitte Gicquel, Ludovic Tailleux

## Abstract

Antibiotics are widely used in the treatment of bacterial infections. Although known for their microbicidal activity, antibiotics may also interfere with the host’s immune system. Here we analyzed the effects of bedaquiline (BDQ), an inhibitor of the mycobacterial ATP synthase, on human macrophages. Genome-wide gene expression analysis revealed that BDQ reprogramed macrophages into potent bactericidal phagocytes. We found that 1,495 genes were differentially expressed in *M. tuberculosis*-infected macrophages incubated with the drug, with an over-representation of genes involved in metabolism, lysosome biogenesis and activation. BDQ treatment triggered a variety of antimicrobial defense mechanisms, including nitric oxide production, phagosome-lysosome fusion, and autophagy. These effects were associated with activation of transcription factor EB (TFEB), involved in the transcription of lysosomal genes, resulting in enhanced intracellular killing of different bacterial species that were naturally insensitive to BDQ. Thus, BDQ could be used as a host-directed therapy against a wide range of bacterial infections.

## Introduction

Antibiotics are commonly used in the treatment of bacterial infections, and, in effectively combating such diseases, have substantially increased human life expectancy. As with most drugs, antibiotic treatment can also alter host metabolism, leading to adverse side-effects, including nausea, hepatotoxicity, skin reactions, and gastrointestinal and neurological disorders. Such side-effects can become critical when antibiotic treatment is long and involves several drugs, as in the treatment of tuberculosis (TB), where 2–28% of patients develop mild liver injury during treatment with first-line drugs (Agal et al., 2005).

Antibiotics can interfere with the immune system, indirectly through the disturbance of the body’s microbiota (Ubeda and Pamer, 2012), or directly by modulating the functions of immune cells. Such interactions may impact treatment efficacy or the susceptibility of the host to concomitant infection. For example, after treatment completion, TB patients are more vulnerable to reactivation and reinfection of the disease, suggesting therapy-related immune impairment (Cox et al., 2008). Drug-sensitive TB can be cured by combining up to 4 antibiotics in a 6-month treatment; specifically, isoniazid (INH), rifampicin (RIF), ethambutol and pyrazinamide (PZA) for 2 months, and INH and RIF for additional 4 months. INH induces apoptosis of activated CD4^+^ T cells in *Mycobacterium tuberculosis* (MTB)-infected mice (Tousif et al., 2014) and leads to a decrease in Th1 cytokine production in household contacts with latent TB under preventive INH therapy (Biraro et al., 2015). RIF has immunomodulatory properties and acts as a mild immunosuppressive agent in psoriasis (Tsankov and Grozdev, 2011). RIF reduces inflammation by inhibiting IκBα degradation, mitogen-activated protein kinase (MAPK) phosphorylation (Bi et al., 2011), and Toll-like receptor 4 signaling (Wang et al., 2013). PZA treatment of MTB-infected human monocytes and mice significantly reduces the release of pro-inflammatory cytokines and chemokines (Manca et al., 2013). It is therefore necessary to understand how antibiotic treatment modulates macrophage (Mφ) functions, and more generally, how it impacts the host immune response.

The world-wide rise in antibiotic resistance is a major threat to global health care. A growing number of bacterial infections, such as pneumonia, salmonellosis, and TB, are becoming harder to treat as the antibiotics used to treat them become less effective. While new antibiotics are being developed and brought to the clinic, their effects on the human immune system are not being studied in-depth. Here, we have investigated the impact of a recently approved anti-TB drug, bedaquiline (BDQ), on the transcriptional responses of human Mφs infected with MTB. Mφs are the primary cell target of MTB, which has evolved several strategies to survive and multiply inside the Mφs phagosome, including prevention of phagosome acidification (Sturgill-Koszycki et al., 1994), inhibition of phagolysosomal fusion (Armstrong and Hart, 1975) and phagosomal rupture (Simeone et al., 2012; van der Wel et al., 2007). They play a central role in the host response to TB pathogenesis, by orchestrating the formation of granulomas, presenting mycobacterial antigens to T cells, and killing the bacillus upon IFN-γ activation (Cambier et al., 2014). BDQ is a diarylquinoline that specifically inhibits a subunit of the bacterial adenosine triphosphate (ATP) synthase, decreasing intracellular ATP levels (Andries et al., 2005; Koul et al., 2007). It has 20,000 times less affinity for human ATP synthase (Haagsma et al., 2009). The most common side effects of BDQ are nausea, joint and chest pain, headache, and arrhythmias (Diacon et al., 2012; Diacon et al., 2014). However, possible interactions between BDQ and the host immune response have not been studied in detail. Understanding the impact of BDQ on the host immune response may help to develop strategies aiming at improving drug efficacy and limiting side-effects, including cytotoxicity, alteration of cell metabolism, and immunomodulation.

## Results

### BDQ modulates the response of MTB-infected Mφs

In order to exclude potential differences due to the MTB bacillary load between treated and untreated cells, we generated a virulent BDQ-resistant strain of *M. tuberculosis* (BDQr-MTB). The selected clone, which carried a Ala63→Pro mutation in subunit c of the ATP synthase (Andries et al., 2005) (*Figure supplement 1A*), had a similar generation time to wild-type bacteria when cultured in 7H9 liquid medium (*Figure supplement 1B*). We also observed no difference in intracellular growth of the mutated and wild-type strains (*Figure supplement 1C*).

We infected human monocyte-derived Mφs from four healthy donors with BDQr-MTB. After 24 h of infection, cells were incubated for an additional 18 h with BDQ at 5 µg/mL, which corresponds to the concentration detected in the plasma of TB patients treated with BDQ (Andries et al., 2005). This concentration did not affect cell viability over an incubation period of 7 days (*Figure supplement 2*). Following treatment, we characterized the genome-wide gene expression profiles of MTB-infected Mφs by RNAseq, with DMSO-treated infected cells serving as a control. The expression of 1,495 genes was affected by BDQ (FDR < 0.05, *Figure 1A and supplementary file 1 and 2*), with 499 being up-regulated and 996 being down-regulated. We classified all 1,495 genes by performing gene-set enrichment analysis using ClueGO cluster analysis (Bindea et al., 2009). The gene set up-regulated by BDQ was significantly enriched for genes associated with glucose/phospholipid metabolism, the lysosome, and autophagy (*Figure 1B*). We observed similar results with uninfected Mφs-treated with BDQ (*Figure supplement 3A-B, supplementary file 3 and 4*), indicating that the effect of BDQ is not dependent on MTB infection.

**Figure 1.**
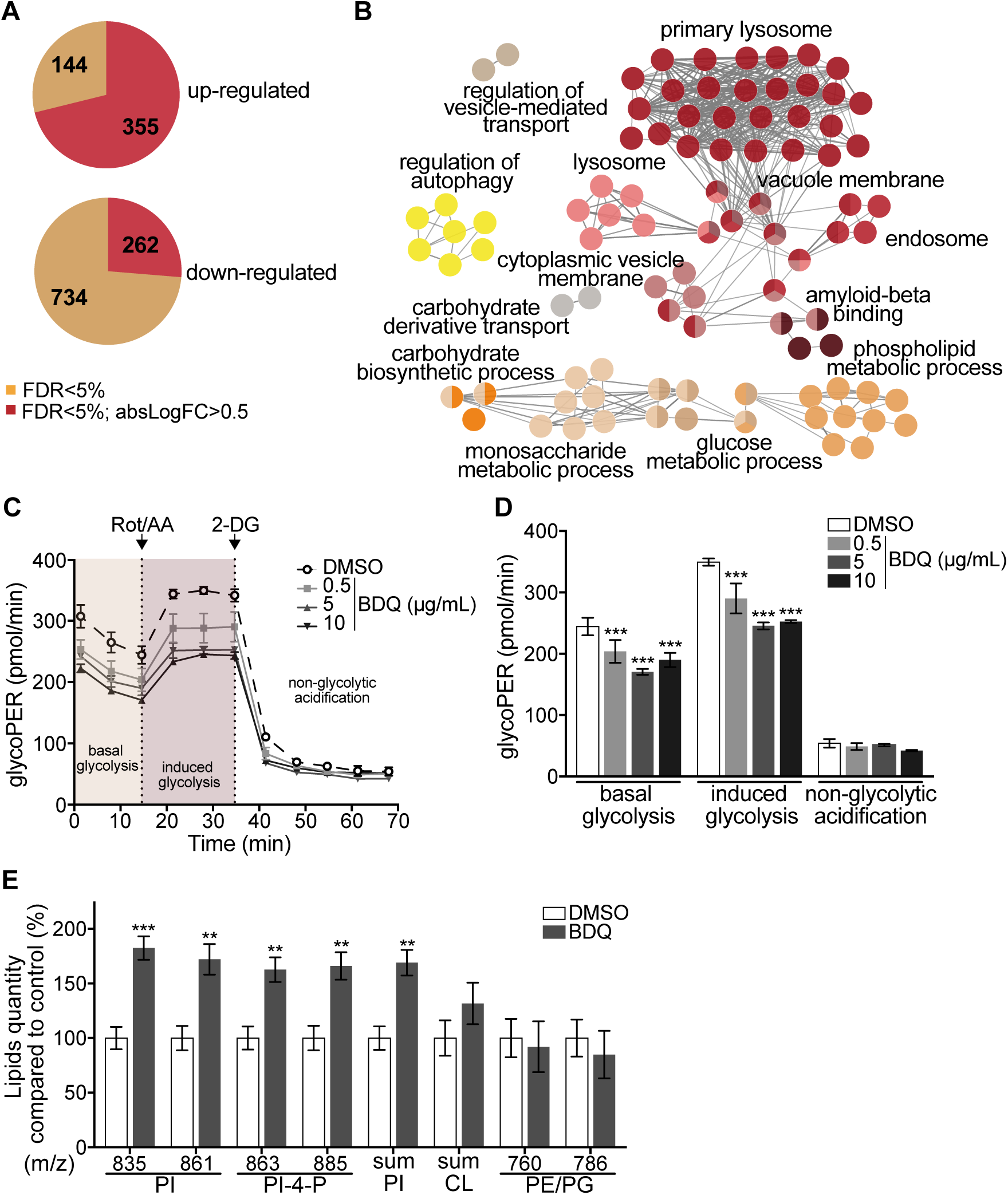
BDQ modulates the response of MTB-infected Mφs. Cells from four donors were infected with BDQ resistant MTB for 24 h and then treated with BDQ (5 µg/mL) for an additional 18 h. Differentially-expressed genes were identified by mRNAseq. See also *Figure supplement 3*. (**A**) Pie chart showing the number of genes regulated by BDQ treatment relative to untreated control. (**B**) Gene ontology enrichment analysis of genes whose expression is up-regulated by BDQ treatment, using the Cytos-cape app ClueGO (FDR<0.05; LogFC>0.5). (**C-D**) The Glycolytic Rate Assay was performed in BDQ-treated Mφs, in the presence of rotenone/antimycin A (Rot/AA) and 2-deoxy-D-glycose (2-DG), inhibitors of the mitochondrial electron transport chain and glycolysis, respectively. (one-way ANOVA test). One representative experiment (of two) is shown. (**E**) Lipid profile of BDQ-treated cells by MALDI-TOF (unpaired two tailed Student’s t test). PI: Phosphotidylinositol; CL: Cardiolipids; PE: Phospha-tidylethanolamine; PG: Phosphatidylglycerol. Numbers correspond to mass-to-charge ratio (m/z). Cells derived from 3 donors were analyzed. Error bars represent the mean ± SD and significant differences between treatments are indicated by an asterisk, in which * p < 0.05, ** p < 0.01, *** p < 0.001.

As metabolic pathways were over-represented in our RNAseq analysis, we investigated if glycolysis is affected by BDQ treatment using the Seahorse Extracellular Flux analyzer. This assay measures the rate of proton accumulation in the extracellular medium during glycolysis (glycoPER) and can discriminate between basal glycolysis, induced glycolytic capacity (by addition of rotenone/antimycin A (Rot/AA), an inhibitor of the mitochondrial electron transport chain), and non-glycolytic acidification (by addition of the glycolytic inhibitor 2-deoxy-D-glycose (2-DG)). After incubation with BDQ, we observed a 30% decrease in basal glycolysis and glycolytic capacity compared to untreated cells (*Figure 1C-D and figure supplement 3C-D*).

We assessed phospholipid metabolism, a pathway also identified in our ClueGO cluster analysis (*Figure 1B*). Like glycolysis, lipid metabolism affects macrophage phenotype and function (Remmerie and Scott, 2018). We analyzed the lipid profile of BDQ-treated cells using MALDI-TOF mass spectrometry. We observed an increase of phosphatidylinositols upon incubation with BDQ (*Figure 1E and figure supplement 3E*). No significant changes were observed in the levels of phosphatidylethanolamines, phosphatidylglycerols, or cardiolipins. Taken together, these data show that BDQ induced a significant metabolic reprogramming of both MTB-infected and resting Mφs.

### BDQ increases Mφ lysosomal activity

Mφs are involved in innate immunity and tissue homeostasis through their detection and elimination of microbes, debris, and dead cells, which occurs in lysosomes (Wynn et al., 2013). Lysosomes are acidic and hydrolytic organelles responsible for the digestion of macromolecules. Recent work has shown that they are also signaling platforms, which respond to nutrient and cellular stress (Lawrence and Zoncu, 2019). Functional annotations based on the KEGG database of the 1,495 genes differentially expressed genes suggested a substantial impact of BDQ treatment on lysosome function (*Figure 1B*). We identified 54 differentially expressed genes, 78% of which were up-regulated, belonging to the lysosomal KEGG term (*Figure 2A*). These genes are involved in lysosome biogenesis, transport and degradation of small molecules, and lysosomal acidification. They included genes coding for components of vacuolar ATPase (V-ATPase), hydrolases, and SLC11A1 (NRAMP1), a divalent transition metal transporter involved in host resistance to pathogens, including MTB (Meilang et al., 2012).

**Figure 2.**
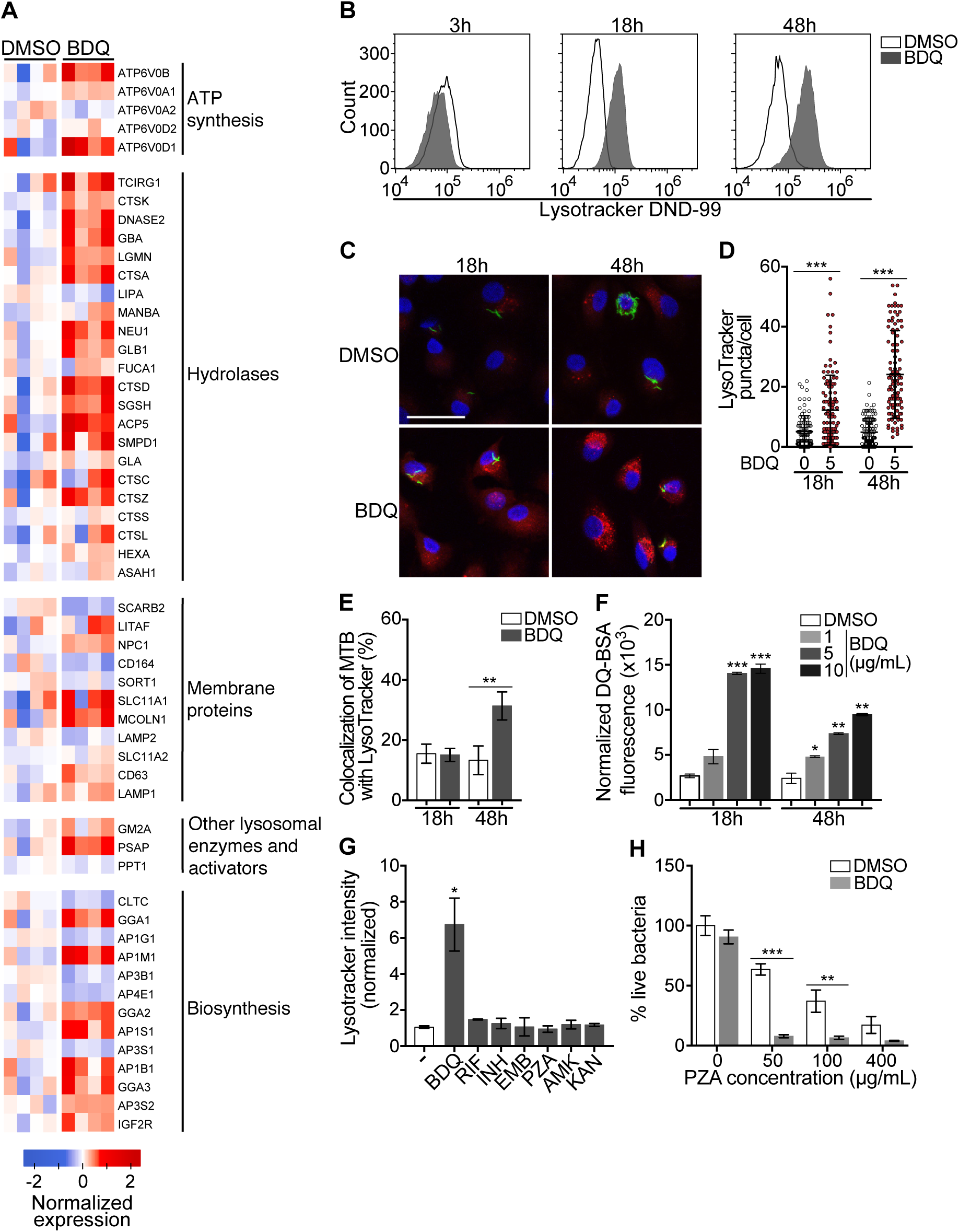
BDQ activates the lysosomal pathway in human MTB-infected Mφs. (**A**) Heatmap showing differential expression of genes included in the Lysosome KEGG category (FDR<0.05%). Each column corresponds to one donor. Data were normalized to determine the log ratio with respect to the median expression of each gene. (**B**) Mφs were infected with BDQr-MTB expressing the GFP protein and incubated with BDQ (5 µg/mL) for 3 h, 18 h and 48 h. Acid organelles were then labeled with 100 nM LysoTracker DND-99 for 1 hour. The fluorescence intensity was quantified by flow cytometry. (**C-E**) Cells were infected with GFP expressing BDQr-MTB (green) and treated with BDQ (5 µg/mL). After 18 h and 48 h of treatment, cells were labelled with LysoTracker (red) and fluorescence was analyzed by confocal microscopy. DAPI (blue) was used to visualize nuclei (scale bar: 10 µm). The quantification of LysoTracker staining and the percentage of LysoTracker-positive MTB phagosomes were performed using Icy software. (**F**) Mφs were activated with heat-killed MTB and treated with BDQ for 18 h and 48 h. Cells were then incubated with DQ-Green BSA. Fluorescence was quantified by flow cytometry. Significant differences between BDQ treatment and control (DMSO) are indicated by an asterisk. (**G**) Mφs were incubated for 48 h with BDQ, rifampicin (RIF, 20 µg/mL), isoniazid (INH, 10 µg/mL), ethambutol (EMB, 20 µg/mL), pyrazinamide (PZA, 200 µg/mL), amikacin (AMK, 20 µg/mL) and kanamycin (KAN, 20 µg/mL), and then stained with LysoTracker. Fluorescence intensity was analyzed by flow cytometry. (**H**) Cells were infected with BDQr-MTB (MOI: 0.5) and treated with BDQ (1 µg/mL) and PZA. After 7 days treatment, cells were lysed and bacteria were enumerated by CFU (counted in triplicate). One representative experiment (of at least three) is shown. Error bars represent the mean ± SD. * p < 0.05, ** p < 0.01, *** p < 0.001.

To validate our transcriptomic data, we incubated BDQ-treated, BDQr-MTB-infected cells with LysoTracker Red DND-99, a red fluorescent probe that labels acidic organelles, and analyzed them using flow cytometry. No differences were observed between control and treatment after 3 h of BDQ treatment (*Figure 2B*). However, at 18 h and 48 h post-treatment, fluorescence intensity was substantially increased in Mφs incubated with BDQ compared to DMSO-treated cells (1.7 and 5.4 times more, respectively). These results were supported by confocal microscopy, which revealed the appearance of numerous acidic compartments upon treatment (*Figure 2C*), up to 5 times more in BDQ-treated Mφs than untreated cells at 48 h post-treatment (p < 0.001, *Figure 2D*). We also observed a large number of MTB phagosomes co-localized with LysoTracker-positive compartments (*Figure 2E*).

As the expression of many genes coding for hydrolases was up-regulated upon BDQ treatment (*Figure 2A*), we tested the effect of the drug on late endosomal/lysosomal proteolytic activity. BDQ-treated Mφs were incubated with DQ-Green BSA, a self-quenched non-fluorescent probe that produces brightly fluorescent peptides following hydrolysis by lysosomal proteases. At 18 h and 48 h post-treatment, we observed a dose-dependent increase in fluorescence intensity upon treatment with BDQ (up to 5.5 times more than untreated cells, p < 0.01, *Figure 2F*). Together, these data demonstrate that BDQ induces biogenesis of competent lysosomes.

Several anti-TB drugs, including INH and PZA, are known to interfere with the degradation and recycling of cellular components (Kim et al., 2012). To test whether other antibiotics might have similar effects to BDQ, we treated cells with amikacin, ethambutol, kanamycin, isoniazid, pyrazinamide, and rifampicin for 48 h and then incubated with LysoTracker (*Figure 2G*). Only BDQ induced an increase in lysosome staining.

The capacity of BDQ to induce acidic compartments may potentiate the efficacy of other drugs, whose activity is pH dependent. In vivo studies have suggested a synergistic interaction between BDQ and PZA (Ibrahim et al., 2007), and it is commonly assumed that a low pH is required for PZA activity against MTB (Zhang and Mitchison, 2003). We thus infected Mφs with BDQr-MTB and treated them with BDQ and PZA. After 7 days of treatment, cells were lysed and bacteria counted. PZA showed moderate bactericidal activity, with 50 µg/mL PZA resulting in a 36% decrease in bacterial numbers compared to untreated cells (*Figure 2H*). We confirmed that the combination of PZA with BDQ was highly bactericidal on MTB, leading to a 83% decrease in colony forming units using 50 µg/mL PZA. This decrease was not a result of an additive effect between the two drugs, as BDQ alone had no antibacterial activity, given that we used a drug-resistant strain of MTB. Thus, the potentiation of PZA activity by BDQ is most likely due to the effect of BDQ on the host cell, and in particular on the increase of lysosomal acidification.

### BDQ induces autophagy activation in Mφs

Given BDQ’s effect on lysosomal acidification we asked whether it promoted lysosome formation. Lysosome biogenesis is linked to the endocytic and autophagic pathways. Autophagy delivers cytoplasmic material and organelles for lysosomal degradation and is implicated in the immune response to microbes (Germic et al., 2019). We therefore tested three inhibitors of the autophagy pathway on BDQ activity: bafilomycin (BAF), which inhibits the V-ATPase; chloroquine (CQ), a lysomotropic agent which prevents endosomal acidification and impairs autophagosome fusion with lysosomes; and 3-methyladenine (3-MA) which blocks autophagosome formation by inhibiting of the type III phosphatidylinositol 3-kinases (PI-3K). We infected Mφs with BDQr-MTB and incubated the cells with BDQ in the presence or absence of the different inhibitors. After 2 days we analyzed LysoTracker staining as a read-out of lysosome activation using flow cytometry and observed that all three inhibitors prevented the increase in staining upon BDQ treatment (*Figure 3A*).

**Figure 3.**
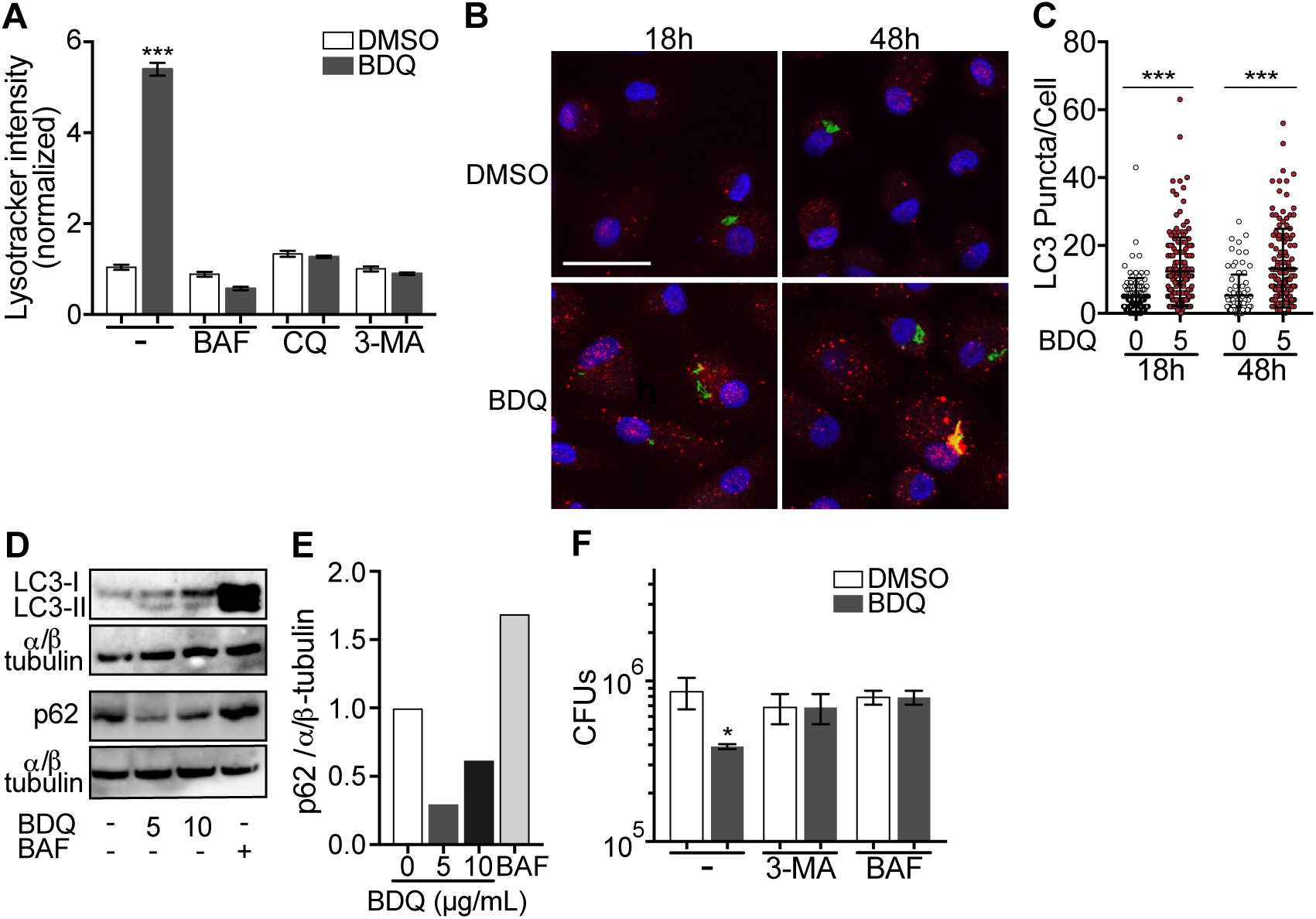
BDQ induced autophagy in MTB-infected Mφs. (**A**) BDQr-MTB infected Mφs were incubated with BDQ (5 µg/mL) and different inhibitors of autophagy; bafilomycin (BAF, 100 nM), chloroquine (CQ, 40 µM) and 3-methyladenine (3-MA, 5 mM). After 48 h, acidic compartments were stained with LysoTracker and fluorescence quantified by flow cytometry. (**B**) Detection by indirect immunofluorescence of LC3 (red) in BDQr-MTB (green) infected Mφs, treated with BDQ for 18 h and 48 h (scale bar: 10 µm). DAPI (blue) was used to visualize nuclei. (**C**) Determination of the number of LC3-positive puncta per cell (one-way ANOVA test). (**D**) Western blot analysis of LC3, p62, and α/ß-tubulin in MTB-infected cells treated with BDQ and BAF. (**E**) Densitometric quantification of p62 staining. (**F**) BDQr-MTB infected Mφs were left untreated or incubated with BDQ, 3-methyladenine (3-MA) and/or bafilomycin (BAF). After 48h, the number of intracellular bacteria was enumerated. One representative experiment (of three) is shown. Error bars represent the mean ± SD. * p < 0.05, ** p < 0.01, *** p < 0.001.

Microtubule-associated protein light-chain 3B (LC3B) is involved in the formation of autophagosomes and autolysosomes. We observed an increase of LC3B puncta per cell at 18 h and 48 h post-BDQ treatment using confocal microscopy (*Figure 3B-C*), which was associated with the detection of lipidated LC3 (LC3-II), the form of LC3 recruited to autophagosomal membranes, and with a decrease in sequestosome 1 (SQSTM1) or p62 levels (*Figure 3D-E*). p62 is a ubiquitin-binding scaffold protein, which is degraded upon autophagy induction, and which is used as a marker of autophagic flux (Liu et al., 2016). Given we have previously observed that some mycobacterial phagosomes colocalized with lysosomes in BDQ-treated cells (*Figure 2E*), we tested whether BDQ promotes MTB killing, independently of its bactericidal activity on MTB by autophagy. BDQ significantly reduced the number of bacteria (measured by CFU) in cells infected with BDQ-resistant MTB. This effect was completely inhibited by the autophagy inhibitors 3-MA and BAF (*Figure 3F*). Overall, these data show BDQ activates the autophagy pathway in human Mφs and this is involved in its anti-TB activity.

### BDQ activates Mφ bactericidal functions

Autophagy plays numerous roles in innate immunity and in host defenses against intracellular pathogens, including MTB (Gutierrez et al., 2004). We thus asked if BDQ conferred protection to bacterial infections naturally resistant to BDQ. To test this hypothesis, we infected Mφs with two different bacterial species: a gram-positive bacterium, *Staphylococcus aureus* and a gram-negative bacterium, *Salmonella* Typhimurium. We confirmed that these two species are resistant to BDQ, even when exposed to high concentration of the drug (20 µg/mL, *Figure 4A*). However, when the cells were incubated with BDQ and then infected with *S. aureus* and *S*. Typhimurium for 24 h, we observed a substantial decrease in bacterial survival rates (*Figure 4B*).

**Figure 4.**
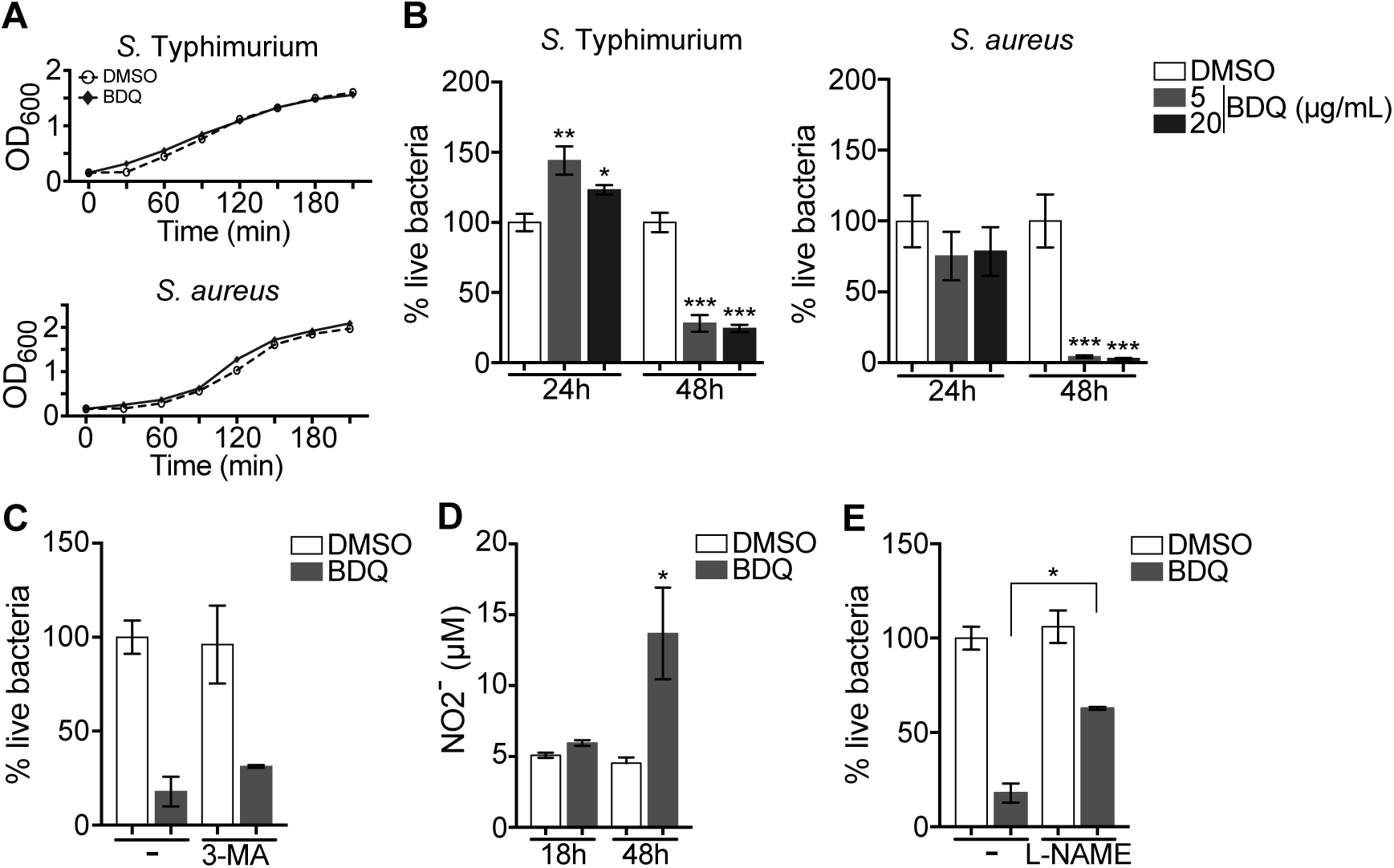
BDQ increases Mφs bactericidal functions. (**A**) Growth of *S*. Typhimurium and *S. aureus* in liquid medium in the presence of BDQ (20 µg/mL). (**B**) Mφs were incubated with BDQ and then infected with *S*. Typhimurium or *S. aureus*. The number of intracellular bacteria was enumerated at 24 h post-infection. (**C**) BDQ-treated Mφs were incubated with 3-MA and then infected with *S. aureus*. The number of bacteria was counted as previously. (**D**) Quantification of NO2-in the supernatant of Mφs incubated with BDQ for 18 h and 48 h. (**E**) Cells were treated as in (*C*), 3-MA was replaced by L-NAME (0.1 mM), an inhibitor of nitric oxide (NO) synthesis. One representative experiment (of three) is shown. Error bars represent the mean ± SD. Unpaired two-tailed Student’s t test was used. * p < 0.05, ** p < 0.01, *** p < 0.001.

To determine if autophagy is involved in this anti-bacterial activity we incubated the infected cells with the autophagy inhibitor, 3-MA, and were unable to revert the Mφs resistance to *S. aureus* infection upon BDQ treatment (*Figure 4C*). Mφs are professional phagocytes, which have evolved a myriad of defense strategies to contain and eradicate bacteria, such as radical formation, phagosome maturation, and metal accumulation (Weiss and Schaible, 2015). Upon incubation with BDQ, we detected an increase in the amount of NO2 -, a stable derivative of NO, in the culture supernatant of Mφs, (*Figure 4D*). When the cells were treated with N(G)-nitro-L-arginine methyl ester (L-NAME), an inhibitor of nitric oxide (NO) synthesis, *S. aureus*-infected cells were unable to effectively control infection upon incubation with BDQ (*Figure 4E*). Thus, our results suggest that BDQ confers innate resistance to bacterial infection through different mechanisms.

### Mitochondrial functions are not affected by BDQ

BDQ affects cardiac electrophysiology by prolonging the QT interval (Diacon et al., 2014) and it has been suggested that BDQ inhibits the cardiac potassium channel protein encoded by the human ether-a-go-go-related gene (hERG) (Therapeutics, 2012). Therefore, to further understand the molecular mechanisms underpinning Mφ activation by BDQ we determined if human monocyte-derived Mφs expressed hERG, but were unable to detect hERG RNA by RT-qPCR (*Figure supplement 4*).

We investigated if BDQ might interfere with other activities of mitochondria. Conflicting reports suggest that BDQ inhibits the mitochondrial ATPase (Fiorillo et al., 2016; Haagsma et al., 2009). We have already shown that there were no significant differences in the amount of cardiolipin, an constituent of inner mitochondrial membranes, between BDQ-treated cells and control cells (*Figure 1E*). We quantified changes in mitochondrial membrane potential using flow cytometry in cells incubated with BDQ or with oligomycin, a positive control, which hyperpolarizes the mitochondrial membrane potential, and stained with TMRM. TMRM is a fluorescent cell-permeant dye that accumulates in active mitochondria with intact membrane potentials. No changes were observed when Mφs were incubated with the BDQ for 6 h, 24 h and 48 h (*Figure 5A*). We obtained similar results when mitochondria were stained with MitoTracker® Red FM whose accumulation in mitochondria is dependent upon membrane potential (*Figure 5B*). We also measured the oxygen consumption rate (OCR), and detected no change in basal respiration, ATP-linked respiration, maximal respiration, and non-mitochondrial respiration in cells treated with BDQ for 24 h and 48 h as compared to untreated cells (*Figure 5C and figure supplement 5*).

Mitochondrial reactive oxygen species (ROS) are involved in the regulation of several physiological and pathological processes, including autophagy (Sena and Chandel, 2012). We thus stained for mitochondrial superoxide using the MitoSOX dye in BDQ-stimulated cells. Again, we saw no difference upon antibiotic treatment (*Figure 5D*). Incubation with the antioxidant glutathione (GSH) or with its precursor N-Acetyl cysteine (NAC), which prevent the formation of mitochondrial ROS and reactive nitrogen species (RNS), did not prevent lysosome activation and the killing of *S. aureus* by BDQ (*Figure 5E*). Based on these results, it is unlikely that BDQ alters mitochondrial function in human Mφs.

### BDQ regulates lysosome activation through TFEB and calcium signaling

Given that BDQ induced a lysosomal gene expression signature in Mφs, we wondered whether BDQ could activate the basic helix-loop-helix transcription factor EB (TFEB). TFEB is a master regulator of autophagy and lysosome biogenesis (Settembre et al., 2011). In resting cells, TFEB is largely cytosolic and inactive, but upon activation, it translocates into the nucleus and activates the transcription of many autophagy and lysosomal genes (Settembre et al., 2011). We therefore analyzed the cellular localization of TFEB, using confocal microscopy. At 18 h post-treatment, TFEB was mainly localized in the nucleus of BDQ-treated cells (*Figure 6A-B*). The activity of TFEB is regulated by phosphorylation on specific amino acid residues, and its activation is mediated by calcineurin, an endogenous serine/threonine phosphatase, through Ca^2+^ release from the lysosome (Medina et al., 2015). In agreement with these studies, we observed an increase in intracellular Ca^2+^ concentration in Mφs treated for 18 h with BDQ (*Figure 6C*), and confirmed that this intracellular calcium accumulation was required for antibiotic-induced TFEB translocation to the nucleus and lysosomal gene expression. Upon treatment with BAPTA, a Ca^2+^ chelator, TFEB remained localized in the cytoplasm of BDQ-treated cells (*Figure 6D*), and we were unable to detect changes in the expression of a panel of lysosomal genes, previously identified as differentially expressed in Mφs incubated with BDQ (*Figure 6E*). The increased bactericidal activity against *S. aureus* was also abrogated in the presence of BAPTA (*Figure 6F*). Collectively, our data indicate that BDQ activates TFEB in Mφs and in this way modulates innate immune resistance to bacterial infection.

**Figure 6.**
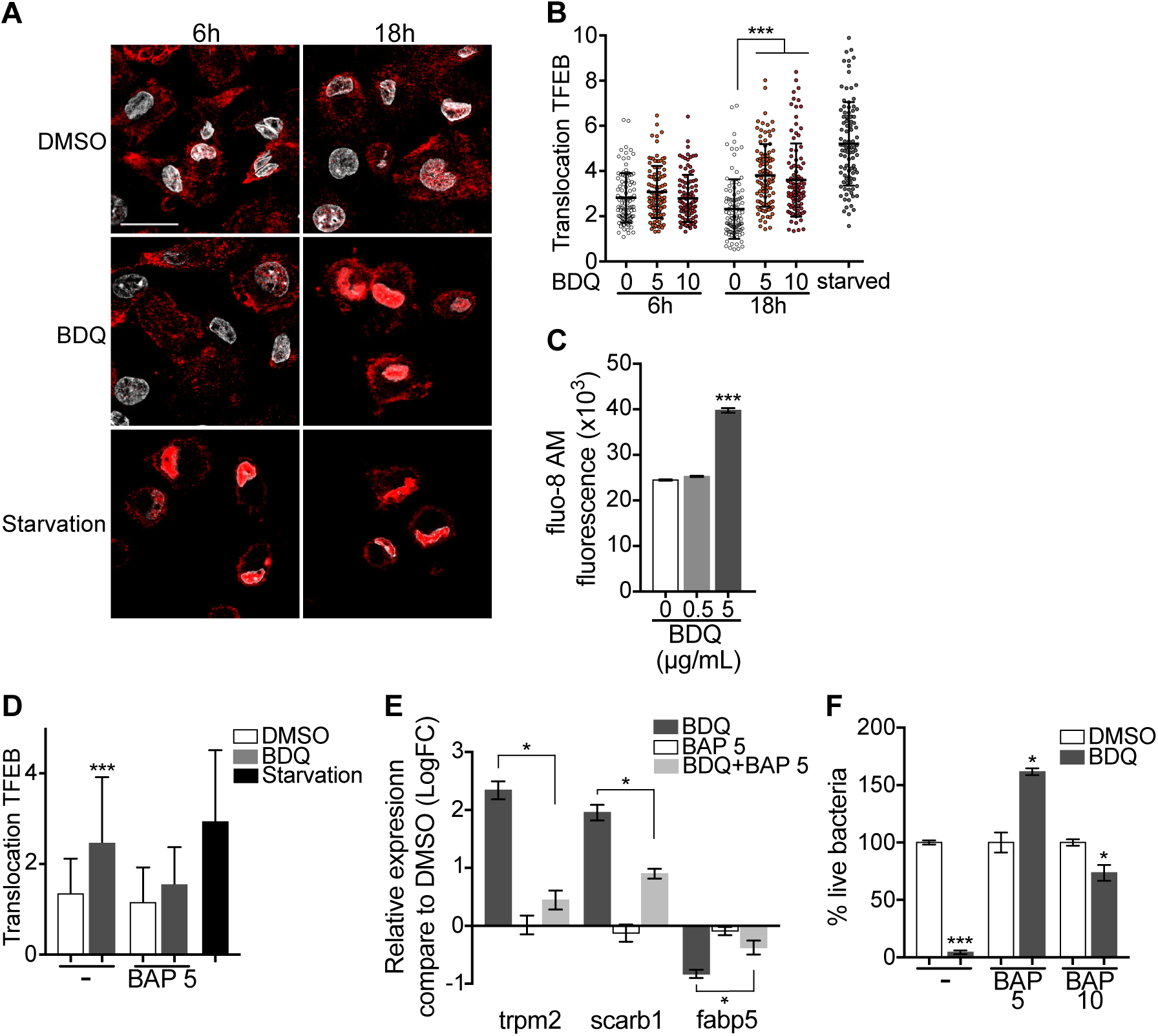
Calcium-dependent activation of TFEB by BDQ. (**A**) Representative fluorescence microscopy images of Mφs treated with BDQ for 6 h and 18 h, or incubated in HBSS medium for 1 h (starvation). Cells were stained with antibody against TFEB (red). DAPI (white) was used to visualize nuclei. Scale bar: 10 µm. (**B**) Ratio between nuclear and cytosolic TFEB fluorescence intensity (n > 100 cells per condition, two-way ANOVA test). (**C**) Mφs were treated with BDQ for 18 h and then loaded with the fluorescent calcium binding dye Fluo-8 AM. After 1 h of incubation, Ca2+ concentration was monitored by FLUOstar Omega. (**D**) Ratio between nuclear and cytosolic TFEB fluorescence intensity in starved cells and in cells treated with BDQ and/or with the intracellular calcium chelator BAPTA (BAP). (n > 100 cells per condition, two-way ANOVA test). (**E**) Relative gene expression measured by RT-qPCR for a panel of differentially expressed lysosomal genes. BDQ treated-Mφs were either left untreated or incubated with BAPTA. Relative expression levels were normalized to the rpl24 gene. (**F**) Mφs were treated with BDQ with or without BAPTA, and then infected with *S. aureus*. After 1 day, the cells were lysed and the number of intracellular bacterial colonies was counted (unpaired two tailed Student’s t test). Error bars represent the mean±SD. was used. * p < 0.05, ** p< 0.01, *** p < 0.001.

## Discussion

The emergence of bacterial strains resistant to antibiotics requires the constant development of new antibiotics, which, beyond their bactericidal activity, may have a significant impact on cellular functions. Here, we have analyzed the effects of the new anti-TB drug BDQ on human Mφs. We found that in addition to its anti-bacterial activity, BDQ induces Mφ cell reprogramming, increasing Mφ bactericidal activity. Gene expression profiling revealed that 1,495 genes were differentially expressed in MTB-infected Mφs incubated with BDQ, with over-representation of genes involved in metabolism, lysosome biogenesis, and acidification. Recent work has highlighted the role of metabolic reprogramming in controlling immunological effector functions, emphasizing the close connection between cell function and metabolism (Wang et al., 2019). In agreement with these results, we observed a substantial increase in both the number of acidic compartments and proteolytic activity of Mφs upon BDQ treatment.

BDQ is a cationic amphiphilic drug, consisting of a hydrophobic ring structure and a hydrophilic side chain with a charged cationic amine group (Diacon et al., 2012). Cationic amphiphilic drugs can accumulate in lysosomes through ion trapping (de Duve et al., 1974). At neutral pH, they passively diffuse across cell and organelle membranes but when they enter the luminal space of acidic compartments such as lysosomes, the amine group ionizes and becomes membrane-impermeable (MacIntyre and Cutler, 1988). Such lysosomotropic compounds usually increase the lysosomal pH and thus decrease lysosomal enzyme activity (Kazmi et al., 2013). However, our results reveal instead that BDQ triggers lysosomal activation, up-regulating the expression of genes coding for hydrolases and for subunits of the lysosomal proton pump v-ATPase. Consistent with these observations, we observed that BDQ-treated cells significantly increase their ability to degrade DQ BSA.

Pre-clinical studies have shown that BDQ may induce phospholipidosis, potentially explaining some of the drug’s observed toxicities (Diacon et al., 2012). Phospholipidosis, which is characterized by the accumulation of phospholipids in lysosomes, resulting in impaired lysosome function, is common upon treatment with cationic amphiphilic compounds (Shayman and Abe, 2013). Various phospholipid species have been described including sphingomyelin, phosphatidylcholine, phosphatidylethanolamine, phosphatidylserine, lysobisphosphatidic acid, and cholesterol (Reasor, 1984; Yamamoto et al., 1971a; Yamamoto et al., 1971b; Yoshikawa, 1991). In BDQ-treated Mφs, we only observed an increase in the amount of phosphatidylinositol and phosphatidylinositol-4-phosphate. The quantity of cardiolipin, phosphatidylethanolamine, and phosphatidylglycerol remained unchanged upon treatment. These observations do not indicate lysosomal dysfunction, but rather a targeted regulation of certain phospholipids by BDQ. In accordance with this idea, 28 genes involved in phospholipid metabolism were differentially expressed in BDQ-treated Mφs. Phosphatidylinositol phosphates regulate many cellular functions, including endosomal trafficking, endoplasmic reticulum (ER) export, autophagy, and phagosome-lysosome fusion (De Matteis et al., 2013; Levin et al., 2017). These phospholipids may thus be involved in the increase of autophagy and mycobacterial phagosome-lysosome fusion upon BDQ treatment. Consistent with this hypothesis, recent work has shown that BDQ accumulates in host cell lipid droplets and is transferred to MTB as the droplets are consumed by the bacteria, enhancing MTB killing (Greenwood et al., 2019).

Lysosomes are both digestive organelles of the endocytic and autophagic pathways and signaling hubs involved in nutrient sensing, cell growth and differentiation, transcriptional regulation, and metabolic homeostasis (Lamming and Bar-Peled, 2019; Lawrence and Zoncu, 2019). In response to nutrients and growth factors, the mechanistic target of the rapamycin complex 1 (mTORC1) is recruited and activated at the lysosomal surface, where it promotes ribosomal biogenesis, translation, and biosynthesis of lipids (Lamming and Bar-Peled, 2019; Lawrence and Zoncu, 2019). mTORC1 binds to and phosphorylates TFEB, resulting in its cytosolic sequestration (Roczniak-Ferguson et al., 2012; Settembre et al., 2012). Upon starvation or lysosomal stress, mTORC1 is released from the lysosomal membrane and becomes inactive (Lamming and Bar-Peled, 2019; Lawrence and Zoncu, 2019). The release of lysosomal Ca^2+^ activates the phosphatase calcineurin, which de-phosphorylates TFEB and promotes its nuclear translocation (Medina et al., 2015). TFEB then binds to CLEAR (coordinated lysosomal expression and regulation) elements within the promoters of genes involved in autophagy and lysosomal biogenesis and activates their expression (Lamming and Bar-Peled, 2019; Lawrence and Zoncu, 2019). We found that TFEB translocates from the cytoplasm to the nucleus in a calcium-dependent manner in BDQ-treated cells, with the concomitant up-regulation of 85 genes containing CLEAR elements 18 h after incubation with the drug.

A striking feature of BDQ-treated Mφs is their capacity to control pathogenic bacterial infection. BDQ enhances Mφ innate defense mechanisms, including induction of anti-microbial effectors such as nitric oxide, phagosome-lysosome fusion, and autophagy. Other anti-TB drugs have been described to regulate autophagy. INH and PZA promote autophagy activation and phagosomal maturation in MTB-infected murine Mφs (Kim et al., 2012) and this process was suggested to be essential for antimycobacterial drug action and for dampening proinflammatory cytokines (Kim et al., 2012). However, a bactericidal effect of INH and PZA could not be excluded as a drug-sensitive MTB strain was used (Kim et al., 2012). In our system, we did not detect increased autophagy in cells treated with INH, which may also be due to differences in the autophagy response in murine and human Mφs. Altogether, we demonstrate that BDQ is able to boost the innate defenses of human cells.

A growing number of pathogenic bacteria are becoming resistant to antibiotics, making their use less effective. In addition to the development of “classical” drugs targeting key factors in bacterial physiology, host-directed therapy (HDT) has emerged as approach that could be used in adjunct with existing or future antibiotics (Machelart et al., 2017). Targeting the host to improve treatment has a number advantages. In particular, HDT is less prone to the development of resistance and it could be used to reduce disease severity and mortality. For example, metformin, an FDA-approved drug for type II diabetes, increases the production of mitochondrial reactive oxygen species and stimulates phagosome-lysosome fusion by activating the 5′-adenosine monophosphate-activated protein kinase (AMPK) (Singhal et al., 2014), and recent studies suggest that metformin provides better outcomes in TB patients, especially those with diabetes mellitus (Yew et al., 2019). Pathogens manipulate host-signaling pathways to subvert innate and adaptive immunity. It might thus be possible to reprogram the host immune system to better control or even kill bacteria. For instance, MTB has developed several strategies to counteract autophagy, including the product of the enhanced intracellular survival (Eis) gene, which limits ROS generation (Shin et al., 2010). Our results clearly show that BDQ can bypass these escape mechanisms and allow more effective control of bacterial infection. We also showed that BDQ potentiates the activity of other anti-TB drugs, independently of its bactericidal activity on MTB. Hence, our work opens new avenues for downstream evaluation of the potential use of BDQ as a potent drug in HDT.

## Materials and methods

### Ethics Statement

Buffy coats were obtained from healthy donors after informed consent. The blood collection protocols were approved by both the French Ministry of Research and a French Ethics Committee. The blood collection was carried out in accordance with these approved protocols by the Etablissement Français du Sang (EFS).

### Φ, MTB and infection

Blood mononuclear cells were isolated from buffy coats by Lymphocytes Separation Medium centrifugation (Eurobio, Les Ulis, France). CD14^+^ monocytes were isolated by positive selection using CD14 microbeads (Miltenyi Biotec, Bergisch Gladbach, Germany) and were allowed to differentiate into Mφs in the presence of granulocyte macrophage colony-stimulating factor (GM-CSF, 20 ng/mL; Miltenyi Biotec) over a 6-day period. To exclude potential differences due to the MTB bacillary load between treated and untreated cells, Mφs were infected with BDQ-resistant MTB strain H37Rv (BDQr-MTB) expressing green-fluorescent protein (GFP). Briefly, exponentially growing MTB carrying the pEGFP plasmid (Tailleux et al., 2003) was plated during 4 weeks on Middlebrook 7H11 agar supplemented with OADC (Becton Dickinson) and containing 0.3 µg/mL BDQ. Some clones were then selected. Resistance to BDQ was confirmed (i) by bacterial culture in Middlebrook 7H9 Broth (Becton Dickinson) supplemented with albumin-dextrose-catalase (ADC, Becton Dickinson) and 0.3 µg/mL BDQ, and (ii) by confirming the mutation in the ATP synthase gene. The *atpE* gene was PCR-amplified using primers (forward: 5-TCGTGTTCATCCTGATCTCCA-3; reverse: 5-GACAATCGCGCTCACTTCAC-3) and the PCR products were sent to Eurofins for sequencing. All the selected mutants carried a mutation in the *atpE* gene as described previously (Andries et al., 2005). Only mutant with similar growth rate (in liquid medium and in Mφs) as the wild type strain has been used for further experiments. Before infection, bacteria were washed and resuspended in 1 mL PBS. Clumps were disassociated by 50 passages through a needle, and then allowed to sediment for 5 min. The density of bacteria in the supernatant was verified by measuring the OD600 and aliquot volumes defined to allow 0.5 bacterium-per-cell infections. After 2 h of incubation at 37 °C, infected cells were thoroughly washed in RPMI 1640 to remove extracellular bacteria and were incubated in fresh medium.

### RNA isolation, library preparation and sequencing

Total RNA from Mφs was extracted using QIAzol lysis reagent (Qiagen, Hilden, Germany) and purified over RNeasy columns (Qiagen). The quality of all samples was assessed with an Agilent 2100 bioanalyzer (Agilent Technologies, Santa Clara, California) to verify RNA integrity. Only samples with good RNA yield and no RNA degradation (ratio of 28S to 18S, >1.7; RNA integrity number, >9) were used for further experiments. cDNA libraries were prepared with the Illumina TruSeq RNA Sample Preparation Kit v2 and were sequenced on an llumina HiSeq 2500 at the CHU Sainte-Justine Integrated Centre for Pediatric Clinical Genomics (Montreal, Canada).

STAR v2.5.0b (Dobin et al., 2013) was used to map RNA-seq reads to the hg38 reference genome and quantify gene expression (option-quantMode GeneCounts) by counting the fragments overlapping the Ensembl genes (GRCh38 v. 83). Differential expression analysis was performed using a generalized linear model with the R Bioconductor package DESeq2 v1.18.1 (Love et al., 2014) on the 12,584 genes with at least one count-per-million (CPM) read in at least four samples. The model formula used in DESeq2 (∼ Donor + Infection + Infection:Donor + Infection:Treatment + Donor:Treatment) contained: the main effects for Donor and Infection, interactions of Donor with Infection and Treatment to adjust for various responses to infection and treatment between donors, and a nested interaction of Infection with Treatment because we were interested in the infection-status-specific treatment effects. The latter was used to extract differentially expressed genes between treated and untreated samples under the infected and uninfected conditions. P-values were adjusted for multiple comparisons using the Benjamini-Hochberg method producing an adjusted P-value or false-discovery rate (FDR).

Gene ontology (GO) enrichment analyses were performed using the Cytoscape app ClueGO (version 2.5.3) (Bindea et al., 2009). The following parameters were used: only pathways with pV ≤ 0.01, Minimum GO level = 3, Maximum GO level = 8, Min GO family > 1, minimum number of genes associated to GO term= 5, and minimum percentage of genes associated to GO term = 8. Enrichment p-values were calculated using a hypergeometric test (p-value < 0.05, Bonferroni corrected).

### Measurement of glycolysis

Measurement of glycolysis was done using the Glycolytic rate assay kit (Seahorse, Agilent Technologies), following the manufacturer’s protocol. Briefly, cells were seeded in Xe96 plates treated with BDQ for 24 h. The cells were then incubated in the assay medium (Seahorse XF Base Medium without phenol, 2mM glutamine, 10 mM glucose, 1 mM pyruvate and 5.0 mM HEPES) at 37°C, during 1 h. Extracellular acidification rate (ECAR, milli pH/min) and oxygen consumption rate (OCR, pmol/min) were measured using the Seahorse Bioscience XFe96 Analyzer.

### Lipidomic

Cells were treated with BDQ during 18 h and them lysed in water during 10 min at 37°C. Samples were heated at 90°C during 40 min in order to inactivate MTB, and were then washed three times to remove salts and contaminants that could preclude the analysis. Prior to mass spectrometry analysis, the 2,5-dihydroxybenzoic acid (Sigma-Aldrich, Saint-Louis, Missouri) matrix was added at a final concentration of 10 mg/mL in a chloroform/methanol mixture at a 90:10 (v/v) ratio; 0.4 µL of a cell solution at a concentration of 2 × 10^5^ to 2 × 10^6^ cells/mL, corresponding to ∼100–1000 cells per well of the MALDI target plate (384 Opti-TOF 123 mm × 84 mm, AB Sciex), and 0.6 µL of the matrix solution were deposited on the MALDI target plate, mixed with a micropipette, and left to dry gently. MALDI-TOF MS analysis was performed on a 4800 Proteomics Analyzer (with TOF-TOF Optics, Applied Biosystems, Foster City, California) using the reflectron mode. Samples were analyzed operating at 20 kV in the negative and positive ion mode. Mass spectrometry data were analyzed using Data Explorer version 4.9 from Applied Biosystems.

### Staining and quantification of acidic compartments

Cells were incubated with LysoTracker DND-99 (100 nM; Thermo Fisher, Waltham, Massachusetts) during 1 h at 37°C. Cells were then fixed with 4% paraformaldehyde at room temperature (RT) for 1 h. Fluorescence was analyzed using a CytoFLEX Flow Cytometer (Beckman Coulter, Brea, California). More than 10,000 events per sample were recorded. The analysis was performed using the FlowJo software.

LysoTracker staining was also analyzed using a Leica TCS SP5 Confocal System. Briefly, cells were washed twice with PBS after incubation with LysoTracker DND-99 (1 µM), fixed with 4% paraformaldehyde for 1 h at RT, stained with DAPI (1 µg/mL, Thermo Fisher) during 10 min mounted on a glass slide using Fluoromount mounting medium (Thermo Fisher). Quantification of LysoTracker staining was performed using Icy software.

### Quantification of lysosomal proteolytic activity

Mφs were activated with heat-killed MTB and treated with BDQ during 18 h or 48 h. Cells were then incubated with DQ-Green BSA (10 µg/mL; Thermo Fisher) for 1 h at 37°C. The hydrolysis of the DQ-Green BSA by lysosomal proteases produces brightly fluorescent peptides. Cells were washed and incubated further in culture medium for 3 h to ensure that DQ BSA had reached the lysosomal compartment. Cells were detached and were fixed with 4% paraformaldehyde and the fluorescence was analyzed using a CytoFLEX Flow Cytometer (Beckman Coulter).

### Determination of bacterial counts

Mφs were lysed in distilled water with 0.1% Triton X-100. MTB was enumerated as previously described (Tailleux et al., 2003) and plated on 7H11. CFUs were scored after three weeks at 37 °C. *S. aureus* and *S*. Typhimurium were plated on Luria-Bertani agar and CFUs were counted after 1 day at 37°C.

### Indirect Immunofluorescence

Mφs (4 × 10^5^ cells/mL) were grown on 12-mm circular coverslips in 24-well tissue culture plates for 24 h in cell culture medium, followed by BDQ treatment. Cells were fixed with 4% paraformaldehyde for 1 h at RT, and were then incubated for 30 min in 1% BSA (Sigma-Aldrich) and 0.075% saponin (Sigma-Aldrich) in PBS, to block nonspecific binding and to permeabilize the cells. Cells were incubated with anti-LC3 (MBL, Woburn, Massachusetts) during 2 h at RT. Alternatively, cells were fixed with cold methanol for 5 min, and were then incubated for 10 min in PBS containing 0.5% saponin. Cells were stained with anti-TFEB (Thermo Fisher) overnight at 4°C. Cells were washed and incubated with Alexa Fluor 555 secondary antibody (Thermo Fisher) for 2 h. Nuclei were stained with DAPI (1 µg/mL) during 10 min. After labeling, coverslips were set in Fluoromount G medium containing 1 µg/ml 4′,6-diamidino-2-phenylindole (DAPI) (SouthernBiotech, Birmingham, Alabama) on microscope slides. Fluorescence was analyzed using Leica TCS SP5. Quantification of TFEB staining was performed using Icy software. LC3B puncta were analysed by confocal microscopy and quantified using ImageJ. Infected cells were manually segmented, thresholded and puncta counted using Analyze Particles. Dot plots represent the mean values of at least 83 cells from 2 donors. Error bars depict the SD.

### Quantitative reverse transcription PCR (RT-qPCR)

Reverse transcription of mRNA to cDNA was done using SuperScript III Reverse Transcriptase (Thermo Fisher) followed by amplification of cDNA using Power SYBR Green PCR Master Mix (Thermo Fisher). The following primers were used: FABP5 (forward: 5-GGAAGGAGAGCACGATAACAAGA-3; reverse: 5’-GGTGGCATTGTTCATGACACA-3), hERG (forward: 5-GGGCTCCATCGAGATCCT-3; reverse: 5-AGGCCTTGCATACAGGTTCA-3), RPL24 (forward: 5-CAAAAGAAAAGAACCCGCCGA-3; reverse: 5-TCGAAACTGGGGAACCATGA-3), SCARB1 (forward: 5-CTTGTTTCTCTCCCATCCTCA-3; reverse: 5-GAGTGTGCCTCCTGGTTAG-3), and TRPM2 (forward: 5’-ACGTGCTCATGGTGGACTTC-3’; reverse: 5’-AGGGTCATAGAAGAGCTGCC-3’). Reactions were performed using a StepOnePlus Real-Time PCR System Thermal Cycling block (Applied Biosystems). The relative gene expression levels were assessed according to the 2^-^ΔCt method (Pfaffl, 2001).

### Western blot analysis

Cells were lysed with RIPA buffer (Thermo Fisher) containing protease inhibitor cocktails (Roche) and stored at −80°C. Protein concentration was determined using the BCA protein assay kit (Thermo Fisher) according to the manufacturer instructions. 20 µg of total protein were loaded on a NUPAGE 4–12% Bis-Tris polyacrylamide gel (Thermo Fisher) and transferred to PVDF membranes (iBlot, Thermo Fisher). The membranes were blocked with TBS-0.1% Tween20, 5% non-fat dry milk for 30 min at RT and incubated overnight with primary antibodies against α-β-Tubulin, p-62 (Cell Signaling) and LC3 (Abcam, Cambridge, United Kingdom). Membranes were washed in TBS-Tween and incubated with secondary HRP-conjugated antibody (GE Healthcare, Chicago, Illinois) at RT for 1 h. Membranes were washed and exposed to SuperSignal West Femto Maximum Sensitivity Substrate (Thermo Fisher). Detection and quantification of band intensities was performed using Azure Imager C400 (Azure Biosystems, Dublin, California) and ImageJ software (version 1.51).

### Infection *S*. *aureus* & *S*. Typhimurium

*S. aureus* and *S*. Typhimurium were grown in Luria-Bertani broth. Bacteria were washed 3 times and resuspended in PBS. The density of bacteria was estimated by measuring the OD_600_. Cells were then infected at a multiplicity of infection of 2:1. After 1 h of infection, cells were extensively washed and incubated for 1 h in culture medium supplemented with gentamicin (100 µg/mL). After washing, cells were cultured with different concentrations of BDQ and gentamicin (5 µg/mL).

### Measurement of nitric oxide

NO was measured by Griess reaction assay (Promega, Madison, Wisconsin) according to the manufacturer’s instructions. Briefly cell culture supernatants were incubated with N-1-napthylethylenediamine dihydrochloride during 10 min followed by additional 10 min with N-1-napthylethylenediamine dihydrochloride. The absorbance was measured at 520 nm.

### Mitochondrial membrane potential

Cells were stained with Image-IT TMRM (10 nM, Thermo Fischer) during 30 min at 37°C or with MitoTracker Deep Red (100nM, Thermo Fisher) during 45 min at 37°C. Cells were washed in PBS and detached from culture plates with 0.05% Trypsin-EDTA. Fluorescence was analyzed using a CytoFLEX Flow Cytometer (Beckman Coulter).

### Measurement of oxygen consumption

The oxygen consumption rate was measured using the XF Cell Mito Stress Test Kit (Seahorse, Agilent Technologies) according to the manufacturer’s protocol. Briefly, cells were seeded in Xe96 plates and treated with BDQ for 24h. The test was performed by adding oligomycin (1 µM), FCCP (1 µM), rotenone and antimycin (0.5 µM) at the indicated time points.

### Mitochondrial ROS assay

Cells were incubated with MitoSOX Red (5 µM, Thermo Fisher) during 10 min at 37°C. Cells were washed in PBS and detached from culture plates with 0.05% Trypsin-EDTA. Fluorescence was analyzed using a CytoFLEX Flow Cytometer (Beckman Coulter).

### Calcium measurement assay

Cells were treated with BDQ for 1 to 18 h, then labeled with Fluo-8 AM (4 µM, Abcam) during 1 h. Cells were washed twice with PBS and fluorescence was analyzed using FLUOstar Omega (BMG Labtech, Ortenberg, Germany).

### Quantification and statistical analysis

Data are expressed as means ± standard deviations (SD). Statistical analyses were performed with Prism software (GraphPad Software Inc.), using the t test and one-way analysis of variance (ANOVA) as indicated in the figure legends. A p value of <0.05 was considered to be significant.

## Data availability

The raw fastq files have been deposited in NCBI’s Gene Expression Omnibus (Edgar et al., 2002) and are accessible through GEO Series accession number GSE133145 (https://www.ncbi.nlm.nih.gov/geo/query/acc.cgi?acc=GSE133145,token:wnkrmgiqnzajxyl).

## Acknowledgments

We thank Olivier Neyrolles for reading the manuscript and helpful discussion. We gratefully acknowledge the UTechS Cytometry and Biomarkers and the UTechS Photonic BioImaging (Imagopole) Citech of Institut Pasteur (Paris, France) as well as the France–BioImaging infrastructure network supported by the French National Research Agency (ANR-10–INSB– 04; Investments for the Future) for support in conducting this study, in particular P.H. Commere for help with flow cytometry. We also thank Charles Privé (CHU Sainte-Justine Integrated Centre for Pediatric Clinical Genomics, Montreal, Canada) for their technical support.

## Additional information

### Additional files

**Supplementary file 1. Up-regulated genes in rBDQ-MTB-infected Mφs upon BDQ treatment**. Related to *Figure 1*. FDR<0.05.

**Supplementary file 2. Down-regulated genes in rBDQ-MTB-infected Mφs upon BDQ treatment**. Related to *Figure 1*. FDR<0.05.

**Supplementary file 3. Up-regulated genes in uninfected Mφs upon BDQ treatment**. Related to *Figure supplement 3*. FDR<0.05.

**Supplementary file 4. Down-regulated genes in uninfected Mφs upon BDQ treatment**. Related to *Figure supplement 3*. FDR<0.05.

---

**Figure supplement 1.**
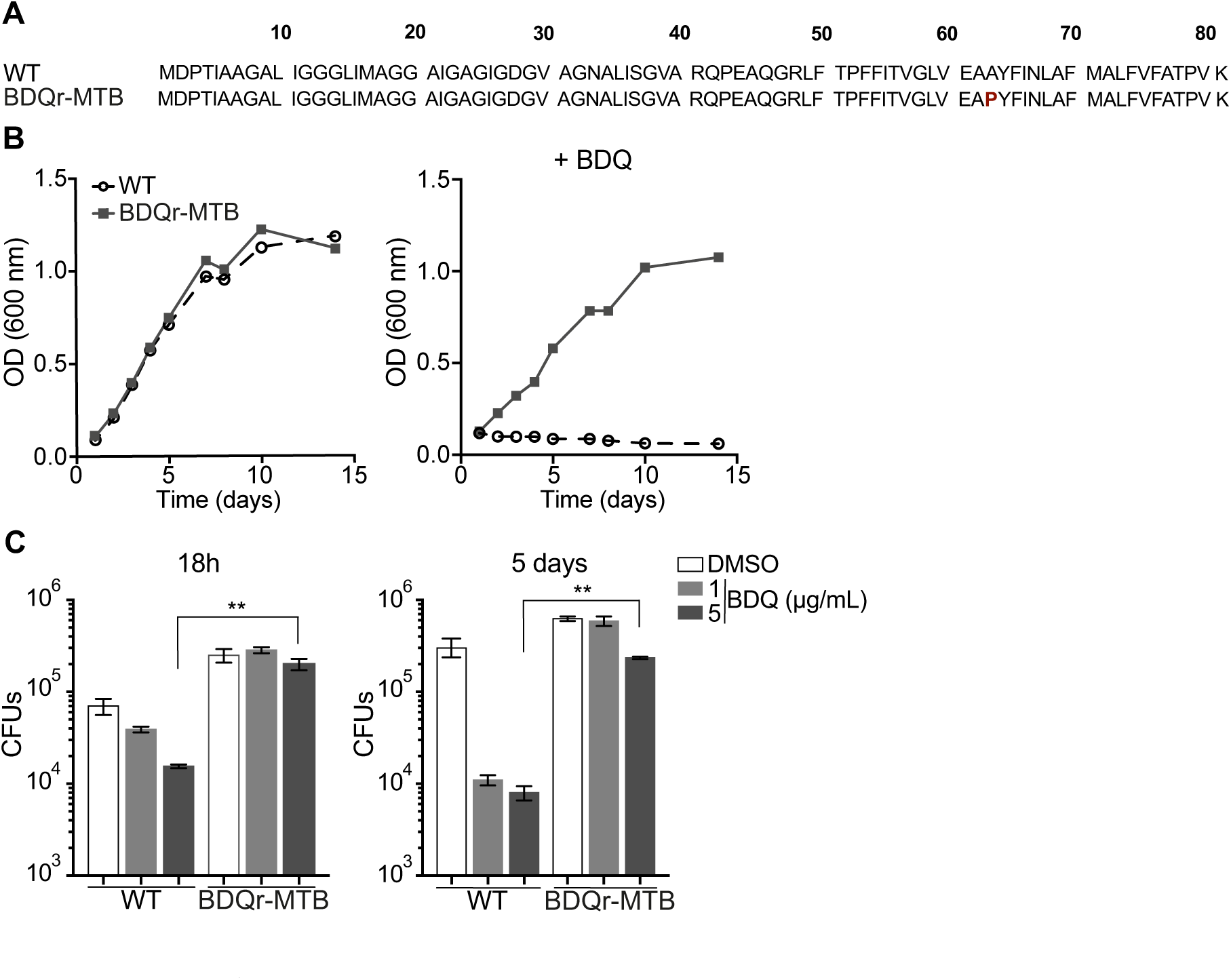
Generation of BDQ resistant MTB strain (BDQr-MTB). (**A**) Amino acid sequence alignment of the ATP synthase c-subunit gene in wild-type (WT) and BDQ-resistant H37Rv strain. The mutation was indicated in red, at position 63. (**B**) Optical density (OD) measurements of bacterial growth of WT and BDQr-MTB. Bacteria were cultured in 7H9 medium supplemented with 10% OADC enrichment with/without BDQ. (**C**) Intracellular growth of wild-type (WT) and BDQ-resistant H37Rv strain. Mφs were infected with the 2 strains and incubated with BDQ. After 18 h and 5 days, the cells were lysed and the number of bacterial colonies was counted. One representative experiment (of three) is shown. Results are means ± SD. ** p < 0.01, unpaired two tailed Student’s t test.

**Figure supplement 2.**
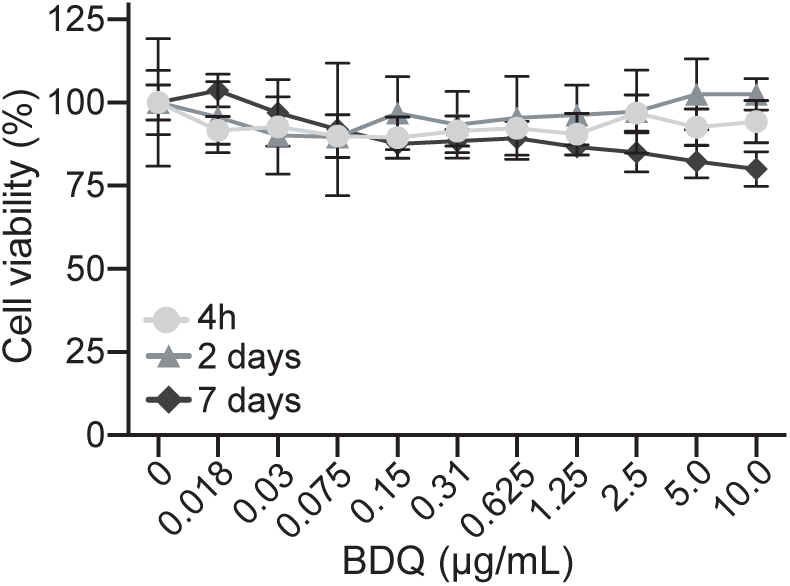
Cell viability assay of Mφs incubated with BDQ. Cells were treated with various concentrations of BDQ. After 4 h, 2 and 7 days, cell viability was evaluated with the MTT assay (Trevigen) according to the manufacturer’s instructions. Results represent the mean ± SD of 3 replicates. One representative experiment (out of three) is shown.

**Figure supplement 3.**
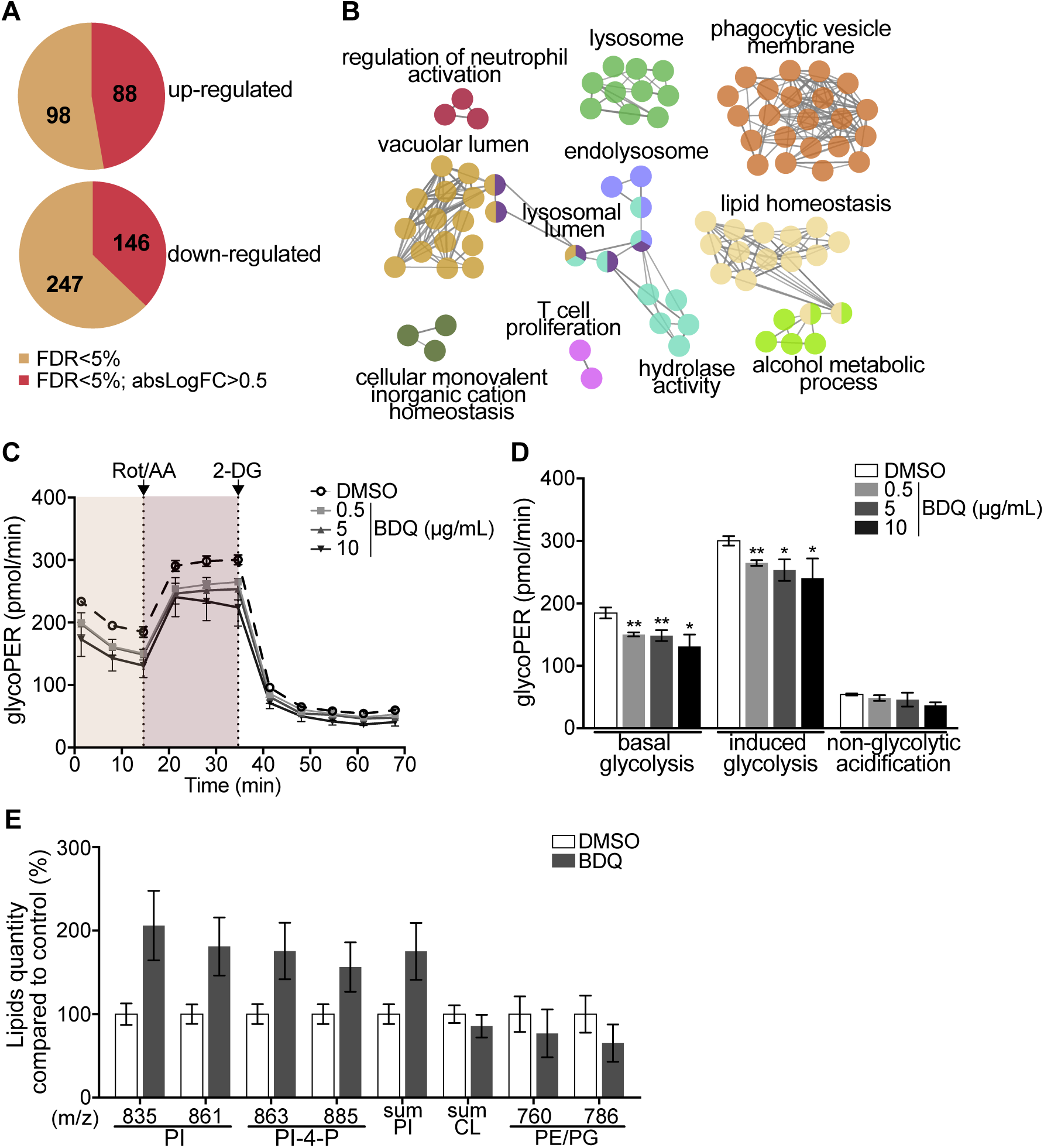
BDQ modulates the response of unactivated Mφs. Related to *Figure 1*. Cells from four individual donors were treated with BDQ (5 µg/mL) for 18 h. The differentially-expressed genes were then identified by mRNAseq. (**A**) Venn diagram showing the number of genes regulated by BDQ treatment relative to untreated controls. (**B**) Gene ontology enrichment analysis of genes whose expression is upregulated by BDQ treatment, using the Cytoscape app ClueGO (FDR<0.05; LogF-C>0.5). (**C-D**) The Glycolytic Rate Assay was performed in Mφs, in the presence of rotenone/antimycin A (Rot/AA) and 2-deoxy-D-glycose (2-DG), respectively inhibitors of mitochondrial electron transport chain and of glycolysis. (one-way ANOVA test). One representative experiment (of two) is shown. (**E**) Lipid profile of cells by MALDI-TOF (unpaired two tailed Student’s t test). PI: Phosphotidylinositol; CL: Cardiolipids; PE: Phosphatidylethanolamine; PG: Phosphatidylglycerol. Numbers correspond to mass-to-charge ratio (m/z). Cells derived from 3 donors were analyzed. Error bars represent the mean ± SD and significant differences between treatments are indicated by an asterisk, in which * p < 0.05, ** p < 0.01, *** p < 0.001.

**Figure supplement 4.**
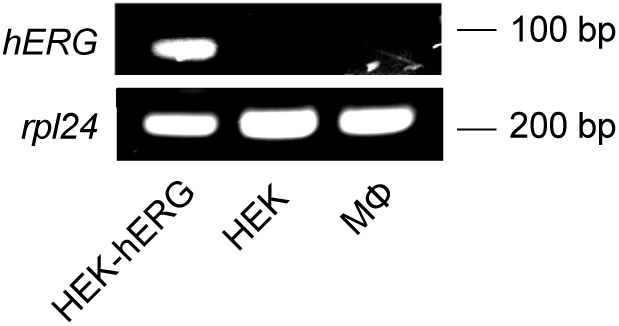
The hERG gene is not expressed in human monocyte-derived Mφs. RT-qPCR was performed in order to detect *hERG* mRNA expression in Mφs, in hERG-transfected and non-transfected HEK293 cells (kind gift from Craig T. January, University of Wisconsin–Madison). *rpl24* was used as control gene.

**Figure supplement 5.**
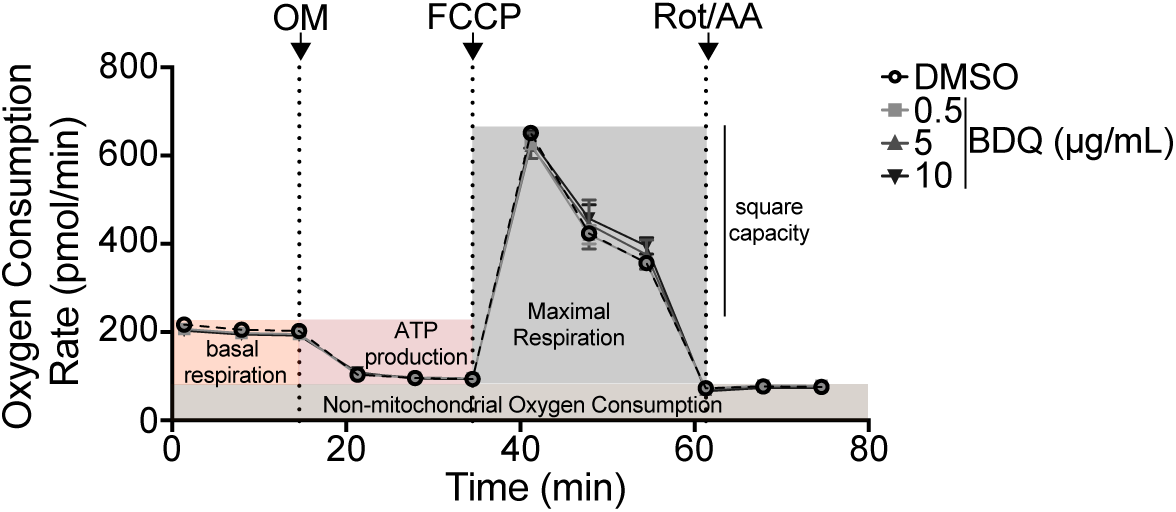
Oxygen consumption rate (OCR) measured by Seahorse extracellular flux assay of cells incubated with BDQ for 48h. Related to *Figure 5*. Basal respiration, ATP production, maximal respiration, respiratory reserve and nonmitochondrial respiration were followed by sequential additions of oligomycin (OM, an inhibitor of the ATPase), the mitochondrial oxidative phosphorylation uncoupler FCCP, and the inhibitors of electron transport antimycin A/rotenone (Rot/AA). Error bars represent the mean ± SD of 3 technical replicates. One representative experiment (out of two) is shown.

## References

Agal, S., Baijal, R., Pramanik, S., Patel, N., Gupte, P., Kamani, P., and Amarapurkar, D. (2005). Monitoring and management of antituberculosis drug induced hepatotoxicity. J Gastroenterol Hepatol. 20(11), 1745–1752. DOI: 10.1111/j.1440-1746.2005.04048.x.

Andries, K., Verhasselt, P., Guillemont, J., Gohlmann, H.W., Neefs, J.M., Winkler, H., Van Gestel, J., Timmerman, P., Zhu, M., Lee, E., et al. (2005). A diarylquinoline drug active on the ATP synthase of Mycobacterium tuberculosis. Science. 307(11), 223–227. DOI: 10.1126/science.1106753.

Armstrong, J.A., and Hart, P.D. (1975). Phagosome-lysosome interactions in cultured macrophages infected with virulent tubercle bacilli. Reversal of the usual nonfusion pattern and observations on bacterial survival. J Exp Med. 142(11), 1–16. DOI: 10.1084/jem.142.1.1.

Bi, W., Zhu, L., Wang, C., Liang, Y., Liu, J., Shi, Q., and Tao, E. (2011). Rifampicin inhibits microglial inflammation and improves neuron survival against inflammation. Brain Res. 1395, 12–20. DOI: 10.1016/j.brainres.2011.04.019.

Bindea, G., Mlecnik, B., Hackl, H., Charoentong, P., Tosolini, M., Kirilovsky, A., Fridman, W.H., Pages, F., Trajanoski, Z., and Galon, J. (2009). ClueGO: a Cytoscape plug-in to decipher functionally grouped gene ontology and pathway annotation networks. Bioinformatics. 25(11), 1091–1093. DOI: 10.1093/bioinformatics/btp101.

Biraro, I.A., Egesa, M., Kimuda, S., Smith, S.G., Toulza, F., Levin, J., Joloba, M., Katamba, A., Cose, S., Dockrell, H.M., et al. (2015). Effect of isoniazid preventive therapy on immune responses to mycobacterium tuberculosis: an open label randomised, controlled, exploratory study. BMC Infect Dis. 15, 438. DOI: 10.1186/s12879-015-1201-8.

Cambier, C.J., Falkow, S., and Ramakrishnan, L. (2014). Host evasion and exploitation schemes of Mycobacterium tuberculosis. Cell. 159(11), 1497–1509. DOI: 10.1016/j.cell.2014.11.024.

Cox, H.S., Morrow, M., and Deutschmann, P.W. (2008). Long term efficacy of DOTS regimens for tuberculosis: systematic review. BMJ. 336(11), 484–487. DOI: 10.1136/bmj.39463.640787.BE.

de Duve, C., de Barsy, T., Poole, B., Trouet, A., Tulkens, P., and Van Hoof, F. (1974). Commentary. Lysosomotropic agents. Biochem Pharmacol. 23(11), 2495–2531.

De Matteis, M.A., Wilson, C., and D’Angelo, G. (2013). Phosphatidylinositol-4-phosphate: the Golgi and beyond. Bioessays. 35(11), 612–622. DOI: 10.1002/bies.201200180.

Diacon, A.H., Donald, P.R., Pym, A., Grobusch, M., Patientia, R.F., Mahanyele, R., Bantubani, N., Narasimooloo, R., De Marez, T., van Heeswijk, R., et al. (2012). Randomized pilot trial of eight weeks of bedaquiline (TMC207) treatment for multidrug-resistant tuberculosis: long-term outcome, tolerability, and effect on emergence of drug resistance. Antimicrob Agents Chemother. 56(11), 3271–3276. DOI: 10.1128/AAC.06126-11.

Diacon, A.H., Pym, A., Grobusch, M.P., de los Rios, J.M., Gotuzzo, E., Vasilyeva, I., Leimane, V., Andries, K., Bakare, N., De Marez, T., et al. (2014). Multidrug-resistant tuberculosis and culture conversion with bedaquiline. N Engl J Med. 371(11), 723–732. DOI: 10.1056/NEJMoa1313865.

Dobin, A., Davis, C.A., Schlesinger, F., Drenkow, J., Zaleski, C., Jha, S., Batut, P., Chaisson, M., and Gingeras, T.R. (2013). STAR: ultrafast universal RNA-seq aligner. Bioinformatics. 29(11), 15–21. DOI: 10.1093/bioinformatics/bts635.

Edgar, R., Domrachev, M., and Lash, A.E. (2002). Gene Expression Omnibus: NCBI gene expression and hybridization array data repository. Nucleic Acids Res. 30(11), 207–210. DOI: 10.1093/nar/30.1.207.

Fiorillo, M., Lamb, R., Tanowitz, H.B., Cappello, A.R., Martinez-Outschoorn, U.E., Sotgia, F., and Lisanti, M.P. (2016). Bedaquiline, an FDA-approved antibiotic, inhibits mitochondrial function and potently blocks the proliferative expansion of stem-like cancer cells (CSCs). Aging (Albany NY). 8(11), 1593–1607. DOI: 10.18632/aging.100983.

Germic, N., Frangez, Z., Yousefi, S., and Simon, H.U. (2019). Regulation of the innate immune system by autophagy: monocytes, macrophages, dendritic cells and antigen presentation. Cell Death Differ. 26(11), 715–727. DOI: 10.1038/s41418-019-0297-6.

Greenwood, D.J., Dos Santos, M.S., Huang, S., Russell, M.R.G., Collinson, L.M., MacRae, J.I., West, A., Jiang, H., and Gutierrez, M.G. (2019). Subcellular antibiotic visualization reveals a dynamic drug reservoir in infected macrophages. Science. 364(11), 1279–1282. DOI: 10.1126/science.aat9689.

Gutierrez, M.G., Master, S.S., Singh, S.B., Taylor, G.A., Colombo, M.I., and Deretic, V. (2004). Autophagy is a defense mechanism inhibiting BCG and Mycobacterium tuberculosis survival in infected macrophages. Cell. 119(11), 753–766. DOI: 10.1016/j.cell.2004.11.038.

Haagsma, A.C., Abdillahi-Ibrahim, R., Wagner, M.J., Krab, K., Vergauwen, K., Guillemont, J., Andries, K., Lill, H., Koul, A., and Bald, D. (2009). Selectivity of TMC207 towards mycobacterial ATP synthase compared with that towards the eukaryotic homologue. Antimicrob Agents Chemother. 53(11), 1290–1292. DOI: 10.1128/AAC.01393-08.

Ibrahim, M., Andries, K., Lounis, N., Chauffour, A., Truffot-Pernot, C., Jarlier, V., and Veziris, N. (2007). Synergistic activity of R207910 combined with pyrazinamide against murine tuberculosis. Antimicrob Agents Chemother. 51(11), 1011–1015. DOI: 10.1128/AAC.00898-06.

Kazmi, F., Hensley, T., Pope, C., Funk, R.S., Loewen, G.J., Buckley, D.B., and Parkinson, A. (2013). Lysosomal sequestration (trapping) of lipophilic amine (cationic amphiphilic) drugs in immortalized human hepatocytes (Fa2N-4 cells). Drug Metab Dispos. 41(11), 897–905. DOI: 10.1124/dmd.112.050054.

Kim, J.J., Lee, H.M., Shin, D.M., Kim, W., Yuk, J.M., Jin, H.S., Lee, S.H., Cha, G.H., Kim, J.M., Lee, Z.W., et al. (2012). Host cell autophagy activated by antibiotics is required for their effective antimycobacterial drug action. Cell Host Microbe. 11(11), 457–468. DOI: 10.1016/j.chom.2012.03.008.

Koul, A., Dendouga, N., Vergauwen, K., Molenberghs, B., Vranckx, L., Willebrords, R., Ristic, Z., Lill, H., Dorange, I., Guillemont, J., et al. (2007). Diarylquinolines target subunit c of mycobacterial ATP synthase. Nat Chem Biol. 3(11), 323–324. DOI: 10.1038/nchembio884.

Lamming, D.W., and Bar-Peled, L. (2019). Lysosome: The metabolic signaling hub. Traffic. 20(11), 27–38. DOI: 10.1111/tra.12617.

Lawrence, R.E., and Zoncu, R. (2019). The lysosome as a cellular centre for signalling, metabolism and quality control. Nat Cell Biol. 21(11), 133–142. DOI: 10.1038/s41556-018-0244-7.

Levin, R., Hammond, G.R., Balla, T., De Camilli, P., Fairn, G.D., and Grinstein, S. (2017). Multiphasic dynamics of phosphatidylinositol 4-phosphate during phagocytosis. Mol Biol Cell. 28(11), 128–140. DOI: 10.1091/mbc.E16-06-0451.

Liu, W.J., Ye, L., Huang, W.F., Guo, L.J., Xu, Z.G., Wu, H.L., Yang, C., and Liu, H.F. (2016). p62 links the autophagy pathway and the ubiqutin-proteasome system upon ubiquitinated protein degradation. Cell Mol Biol Lett. 21, 29. DOI: 10.1186/s11658-016-0031-z.

Love, M.I., Huber, W., and Anders, S. (2014). Moderated estimation of fold change and dispersion for RNA-seq data with DESeq2. Genome Biol. 15(11), 550. DOI: 10.1186/s13059-014-0550-8.

Machelart, A., Song, O.R., Hoffmann, E., and Brodin, P. (2017). Host-directed therapies offer novel opportunities for the fight against tuberculosis. Drug Discov Today. 22(11), 1250–1257. DOI: 10.1016/j.drudis.2017.05.005.

MacIntyre, A.C., and Cutler, D.J. (1988). Role of lysosomes in hepatic accumulation of chloroquine. J Pharm Sci. 77(11), 196–199.

Manca, C., Koo, M.S., Peixoto, B., Fallows, D., Kaplan, G., and Subbian, S. (2013). Host targeted activity of pyrazinamide in Mycobacterium tuberculosis infection. PLoS One. 8(11), e74082. DOI: 10.1371/journal.pone.0074082.

Medina, D.L., Di Paola, S., Peluso, I., Armani, A., De Stefani, D., Venditti, R., Montefusco, S., Scotto-Rosato, A., Prezioso, C., Forrester, A., et al. (2015). Lysosomal calcium signalling regulates autophagy through calcineurin and TFEB. Nat Cell Biol. 17(11), 288–299. DOI: 10.1038/ncb3114.

Meilang, Q., Zhang, Y., Zhang, J., Zhao, Y., Tian, C., Huang, J., and Fan, H. (2012). Polymorphisms in the SLC11A1 gene and tuberculosis risk: a meta-analysis update. Int J Tuberc Lung Dis. 16(11), 437–446. DOI: 10.5588/ijtld.10.0743.

Pfaffl, M.W. (2001). A new mathematical model for relative quantification in real-time RT- PCR. Nucleic Acids Res. 29(11), e45. DOI: 10.1093/nar/29.9.e45.

Reasor, M.J. (1984). Phospholipidosis in the alveolar macrophage induced by cationic amphiphilic drugs. Fed Proc. 43(11), 2578–2581.

Remmerie, A., and Scott, C.L. (2018). Macrophages and lipid metabolism. Cell Immunol. 330, 27–42. DOI: 10.1016/j.cellimm.2018.01.020.

Roczniak-Ferguson, A., Petit, C.S., Froehlich, F., Qian, S., Ky, J., Angarola, B., Walther, T.C., and Ferguson, S.M. (2012). The transcription factor TFEB links mTORC1 signaling to transcriptional control of lysosome homeostasis. Sci Signal. 5(11), ra42. DOI: 10.1126/scisignal.2002790.

Sena, L.A., and Chandel, N.S. (2012). Physiological roles of mitochondrial reactive oxygen species. Mol Cell. 48(11), 158–167. DOI: 10.1016/j.molcel.2012.09.025.

Settembre, C., Di Malta, C., Polito, V.A., Garcia Arencibia, M., Vetrini, F., Erdin, S., Erdin, S.U., Huynh, T., Medina, D., Colella, P., et al. (2011). TFEB links autophagy to lysosomal biogenesis. Science. 332(11), 1429–1433. DOI: 10.1126/science.1204592.

Settembre, C., Zoncu, R., Medina, D.L., Vetrini, F., Erdin, S., Erdin, S., Huynh, T., Ferron, M., Karsenty, G., Vellard, M.C., et al. (2012). A lysosome-to-nucleus signalling mechanism senses and regulates the lysosome via mTOR and TFEB. EMBO J. 31(11), 1095–1108. DOI: 10.1038/emboj.2012.32.

Shayman, J.A., and Abe, A. (2013). Drug induced phospholipidosis: an acquired lysosomal storage disorder. Biochim Biophys Acta. 1831(11), 602–611. DOI: 10.1016/j.bbalip.2012.08.013.

Shin, D.M., Jeon, B.Y., Lee, H.M., Jin, H.S., Yuk, J.M., Song, C.H., Lee, S.H., Lee, Z.W., Cho, S.N., Kim, J.M., et al. (2010). Mycobacterium tuberculosis eis regulates autophagy, inflammation, and cell death through redox-dependent signaling. PLoS Pathog. 6(11), e1001230. DOI: 10.1371/journal.ppat.1001230.

Simeone, R., Bobard, A., Lippmann, J., Bitter, W., Majlessi, L., Brosch, R., and Enninga, J. (2012). Phagosomal rupture by Mycobacterium tuberculosis results in toxicity and host cell death. PLoS Pathog. 8(11), e1002507. DOI: 10.1371/journal.ppat.1002507.

Singhal, A., Jie, L., Kumar, P., Hong, G.S., Leow, M.K., Paleja, B., Tsenova, L., Kurepina, N., Chen, J., Zolezzi, F., et al. (2014). Metformin as adjunct antituberculosis therapy. Sci Transl Med. 6(11), 263ra159. DOI: 10.1126/scitranslmed.3009885.

Sturgill-Koszycki, S., Schlesinger, P.H., Chakraborty, P., Haddix, P.L., Collins, H.L., Fok, A.K., Allen, R.D., Gluck, S.L., Heuser, J., and Russell, D.G. (1994). Lack of acidification in Mycobacterium phagosomes produced by exclusion of the vesicular proton-ATPase. Science. 263(11), 678–681. DOI: 10.1126/science.8303277.

Tailleux, L., Neyrolles, O., Honore-Bouakline, S., Perret, E., Sanchez, F., Abastado, J.P., Lagrange, P.H., Gluckman, J.C., Rosenzwajg, M., and Herrmann, J.L. (2003). Constrained intracellular survival of Mycobacterium tuberculosis in human dendritic cells. J Immunol. 170(11), 1939–1948. DOI: 10.4049/jimmunol.170.4.1939.

Therapeutics, J. (2012). Sirturo (bedaquiline). US Food and Drug Administration, center for drug evaluation and research. https://www.accessdata.fda.gov/drugsatfda_docs/nda/2012/204384Orig1s000PharmR.pdf

Tousif, S., Singh, D.K., Ahmad, S., Moodley, P., Bhattacharyya, M., Van Kaer, L., and Das, G. (2014). Isoniazid induces apoptosis of activated CD4+ T cells: implications for post-therapy tuberculosis reactivation and reinfection. J Biol Chem. 289(11), 30190–30195. DOI: 10.1074/jbc.C114.598946.

Tsankov, N., and Grozdev, I. (2011). Rifampicin--a mild immunosuppressive agent for psoriasis. J Dermatolog Treat. 22(11), 62–64. DOI: 10.3109/09546630903496975.

Ubeda, C., and Pamer, E.G. (2012). Antibiotics, microbiota, and immune defense. Trends Immunol. 33(11), 459–466. DOI: 10.1016/j.it.2012.05.003.

van der Wel, N., Hava, D., Houben, D., Fluitsma, D., van Zon, M., Pierson, J., Brenner, M., and Peters, P.J. (2007). M. tuberculosis and M. leprae translocate from the phagolysosome to the cytosol in myeloid cells. Cell. 129(11), 1287–1298. DOI: 10.1016/j.cell.2007.05.059.

Wang, A., Luan, H.H., and Medzhitov, R. (2019). An evolutionary perspective on immunometabolism. Science. 363(6423). DOI: 10.1126/science.aar3932.

Wang, X., Grace, P.M., Pham, M.N., Cheng, K., Strand, K.A., Smith, C., Li, J., Watkins, L.R., and Yin, H. (2013). Rifampin inhibits Toll-like receptor 4 signaling by targeting myeloid differentiation protein 2 and attenuates neuropathic pain. FASEB J. 27(11), 2713–2722. DOI: 10.1096/fj.12-222992.

Weiss, G., and Schaible, U.E. (2015). Macrophage defense mechanisms against intracellular bacteria. Immunol Rev. 264(11), 182–203. DOI: 10.1111/imr.12266.

Wynn, T.A., Chawla, A., and Pollard, J.W. (2013). Macrophage biology in development, homeostasis and disease. Nature. 496(11), 445–455. DOI: 10.1038/nature12034.

Yamamoto, A., Adachi, S., Ishikawa, K., Yokomura, T., and Kitani, T. (1971a). Studies on drug-induced lipidosis. 3. Lipid composition of the liver and some other tissues in clinical cases of “Niemann-Pick-like syndrome” induced by 4,4’-diethylaminoethoxyhexestrol. J Biochem. 70(11), 775–784. DOI: 10.1093/oxfordjournals.jbchem.a129695.

Yamamoto, A., Adachi, S., Kitani, T., Shinji, Y., and Seki, K. (1971b). Drug-induced lipidosis in human cases and in animal experiments. Accumulation of an acidic glycerophospholipid. J Biochem. 69(11), 613–615.

Yew, W.W., Chang, K.C., Chan, D.P., and Zhang, Y. (2019). Metformin as a host-directed therapeutic in tuberculosis: Is there a promise? Tuberculosis (Edinb). 115, 76–80. DOI: 10.1016/j.tube.2019.02.004.

Yoshikawa, H. (1991). Effects of drugs on cholesterol esterification in normal and Niemann- Pick type C fibroblasts: AY-9944, other cationic amphiphilic drugs and DMSO. Brain Dev. 13(11), 115–120.

Zhang, Y., and Mitchison, D. (2003). The curious characteristics of pyrazinamide: a review. Int J Tuberc Lung Dis. 7(11), 6–21.

